# Joint Estimation of Pedigrees and Effective Population Size Using Markov Chain Monte Carlo

**DOI:** 10.1101/492678

**Authors:** Amy Ko, Rasmus Nielsen

## Abstract

Pedigrees provide the genealogical relationships among individuals at a fine resolution and serve an important function in many areas of genetic studies. One such use of pedigree information is in the estimation of the short-term effective population size (*N*_*e*_), which is of great relevance in fields such as conservation genetics. Despite the usefulness of pedigrees, however, they are often an unknown parameter and must be inferred from genetic data. In this study, we present a Bayesian method to jointly estimate pedigrees and *N*_*e*_ from genetic markers using Markov Chain Monte Carlo. Our method supports analysis of a large number of markers and individuals with the use of a composite likelihood, which significantly increases computational efficiency. We show on simulated data that our method is able to jointly estimate relationships up to first cousins and *N*_*e*_ with high accuracy. We also apply the method on a real dataset of house sparrows to reconstruct their previously unreported pedigree.

## Introduction

Pedigrees are fundamental in many areas of genetic studies. Pedigree structure can be used to study the social organization of a population, such as the degree of polygamy and the offspring distribution among mothers and fathers (Blouin 2003). In conservation genetics, pedigrees provide a way to design an appropriate breeding scheme by preventing inbreeding between close relatives. Other uses of pedigree information include estimating heritability of quantitative traits (Vinkhuyzen *et al.* 2013); controlling for cryptic relatedness in association studies (Voight and Pritchard 2005; Eu-ahsunthornwattana *et al.* 2014); and pedigree-based association studies (Ott *et al.* 2011). Furthermore, the genealogical history embedded in pedigrees can be used to estimate demographic parameters for the recent past, such as the short-term effective population size (*N*_*e*_) (Wang 2009). However, most population genetic models are based on King-man’s coalescent (Kingman 1982a,b,c), which is a poor approximation of the genealogical process for time frames shorter than *log*_2_*N*, where *N* is the population size (Wakeley *et al.* 2012, 2016). Pedigrees, which provide a finer resolution on the genealogical history of the samples than the coalescent, may therefore be more appropriate to use for estimating demographic parameters of the very recent past.

Despite the importance of pedigrees in genetic analyses, they are often a missing parameter. To address this problem, many methods have been developed to estimate pedigrees from genetic data. Existing methods fall broadly into two categories: those that estimate pairwise relationships only (Thompson 1975; McPeek and Sun 2000; Smith *et al.* 2001; Sun *et al.* 2001; Milligan 2003; Sun and Dimitromanolakis 2014) and those that aim to reconstruct the entire pedigree (Thomas and Hill 2000; Almudevar 2003; Wang 2004; Hadfield *et al.* 2006; Gasbarra *et al.* 2007; Cowell 2009; Riester *et al.* 2009; Wang and Santure 2009; Kirkpatrick *et al.* 2011; Almudevar and Anderson 2012; Wang 2012; Cowell 2013; He *et al.* 2013; Cussens *et al.* 2013; Staples *et al.* 2014; Anderson and Ng 2016; Staples *et al.* 2016; Ko and Nielsen 2017; Ramstetter *et al.* 2018). Although pairwise methods are computationally fast, estimated pairwise relationships do not necessarily translate to the correct pedigree, as piecing together pairwise relationships may not produce a valid pedigree. Furthermore, because the coefficient of variation in genome sharing between two individuals becomes larger as the relationship becomes more distant (Hill and Weir 2011), distinguishing competing relationships from each other becomes increasingly difficult. Methods that estimate the entire pedigree have an advantage in this regard. Several studies have shown that the accuracy of pairwise relationship inference can be improved by considering all relationships in the sample simultaneously and resolving uncertain relationships in the context of other individuals (Staples *et al.* 2014; Ko and Nielsen 2017; Ramstetter *et al.* 2018). Furthermore, the estimated pedigree is valid by construction, which can then be used to study population parameters of interest, such as the variance in offspring distribution.

Existing pedigree reconstruction methods, however, are limited in their scope due to the inherent difficulty in pedigree inference. First, the likelihood computation of a pedigree is expensive, as it scales exponentially either in the number of individuals in the pedigree (Lander and Green 1987) or in the number of markers analyzed (Elston and Stewart 1971). Second, the number of possible pedigrees for a given number of individuals is enormous, much greater than the number of phylogenetic trees (Steel and Hein 2006; Thatte and Steel 2008), which makes exploring the pedigree space computationally challenging for even a small number of individuals.

In this study, we present a pedigree inference method that addresses the difficulties of pedigree inference. First, we use the composite likelihood developed in (Ko and Nielsen 2017) to make the likelihood computation efficient for a large number of markers and individuals. Second, we use Markov Chain Monte Carlo (MCMC) (Hastings 1970) to sample pedigrees from high probability regions, circumventing the need to enumerate all possible pedigrees. Our method is different in several important ways from previous methods such as (Wang 2012; Staples *et al.* 2014; Ko and Nielsen 2017) that also use composite likelihoods and sampling algorithms to explore the pedigree space. The previous methods take a maximum likelihood approach and produce a list of pedigrees with highest likelihoods, and does not provide a principled way to compute the uncertainty of the estimated pedigrees. In contrast, our method casts the problem in a Bayesian framework and estimates the posterior probability distribution of the parameters, which in turn quantifies the uncertainty in parameter estimation.

Furthermore, by assigning a prior, which is a function of population parameters that govern the mating behavior of the population, to the pedigrees, we can estimate these parameters jointly with the pedigree. In particular, we focus on estimating the short-term *N*_*e*_, a key parameter quantifying the level of genetic drift and inbreeding in the current population. Various approaches have been developed for estimating the short-term *N*_*e*_, including methods based on relatedness, heterozygosity excess, linkage disequilibrium, or changes in allele frequency over time (Wang *et al.* 2016). Our pedigree-based approach for estimating *N*_*e*_ is most closely related to the estimation method based on the frequency of siblings in a sample by (Wang 2009), which was shown to be more accurate and robust than other approaches.

In our method, we assume that all individuals belong to a single generation and infer pedigrees going up to two generations back in time (i.e. up to first cousins). Furthermore, we assume that the population is outbred with non-overlapping generations and the pedigrees do not contain cycles other than those caused by full sibling relationships. We validate our method on simulated data and show that it can estimate relationships and *N*_*e*_ accurately. Furthermore, we apply our method on a real dataset containing a sample of house sparrows to reconstruct their previously unreported pedigree.

## Materials and Methods

### Bayesian Inference of Pedigrees and Mating Parameters

Our method aims to estimate the joint posterior distribution of pedigrees and mating parameters. Let *n* be the sample size, *H* the pedigree of the sample, *θ* the set of mating parameters for the population, and *X* = (*X*_1_, …, *X*_*n*_) the set of genotype vectors for the *n* individuals. Then the joint posterior probability of *H* and *θ* can be written as

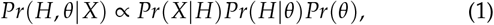

where *Pr*(*X|H*) is the likelihood of the pedigree, *Pr*(*H|θ*) is the prior for the pedigree under a mating model parameterized by *θ*, and *Pr*(*θ*) is the hyperprior on the mating parameters. We describe below how to compute each of these component terms in more detail.

#### Composite Likelihood

As discussed in Introduction, computing the likelihood of a pedigree, *Pr*(*X|H*), is computationally prohibitive for even a moderately large set of markers or individuals. We therefore approximate the likelihood with the composite likelihood introduced in (Ko and Nielsen 2017) to make the computation more efficient. The composite likelihood is based on the marginal pairwise likelihoods, which we describe briefly here.

The composite likelihood of pedigree *H* containing *k* sampled individuals is given by

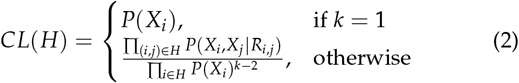

where *R*_*i*,*j*_ is the relationship between individuals *i* and *j* induced by pedigree *H*. If the pedigree contains a single individual (i.e. *k* = 1), then the composite likelihood is simply the probability of observing the individual’s genotypes. For *k* > 1, the composite likelihood is the product of the pairwise likelihoods, scaled by the marginal likelihoods of the individuals. That is, since each individual appears *k* − 1 times in the numerator, we divide the numerator by the marginal likelihood of each individual *k* − 2 times. A previous study by (Ko and Nielsen 2017) showed that the composite likelihood scales similarly to the full likelihood on simulated data and has sensible asymptotic properties, making it a good approximation for the full likelihood.

We pre-compute and store in memory the pairwise likelihoods *Pr*(*X*_*i*_, *X*_*j*_*|R*_*i,j*_) for each pair (*i, j*) for a specified set of pairwise relationships. For pedigrees going up to two generations back in time, this set includes full siblings, half siblings, full first cousins, half first cousins, and unrelated. The pairwise likelihoods can be computed efficiently using the method described in (Weir *et al.* 2006) for unlinked markers or by (Albrechtsen *et al.* 2009) for linked markers. The pairwise likelihoods can then be accessed from memory to compute the composite likelihood efficiently.

#### Prior

For the prior on the pedigrees, *Pr*(*H|θ*), we used a modified version of the mating model introduced in (Gasbarra *et al.* 2005). The model is defined by three parameters: *α, β*, and *N*, which we describe in more detail below.

The probability of a pedigree under this mating model is most naturally described by the procedure by which each child stochastically chooses its mother and father. We assume a homogeneous population of constant size *N* with non-overlapping generations and equal proportions of males and females (i.e. *N*/2 males and *N*/2 females). Let *n* be the number of children in the current generation. One by one, each child chooses a parental pair (*f, m*) where *f* ∈ {1, 2, …, *N*/2} and *m* ∈ {1, 2, …, *N*/2}.

Let *C*_*f*_ (*k*) be the number of children that mother *f* has after the first *k* children have chosen their parents. Then the probability that the (*k* + 1)^*th*^ child chooses mother *f* is given by

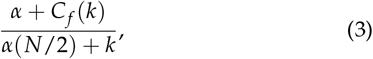

where *α* is a parameter that controls the offspring distribution among mothers in the population. A small value of *α* corresponds to the mating model where a few mothers have many offspring, whereas a large value of *α* corresponds to the model where children are distributed more evenly among all mothers.

After selecting mother *f*, the child chooses a father next. Let *C*_*fm*_(*k*) be the number of children that parental pair (*f, m*) has after the first *k* children have chosen their parents. Then the probability of the (*k* + 1)^*th*^ child choosing father *m* is given by

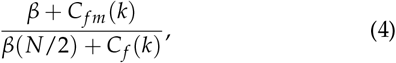

where *β* is a parameter that governs the degree of polygamy of fathers. If *β* is small, then the child is more likely to choose father *m* if the father already shares other offspring with the child’s mother, *f* (i.e. parental pairs tend to stay monogamous). On the other hand, *β* = ∞ corresponds to the case where the child chooses a father at random (i.e. random mating model).

After all *n* children in the current generation have chosen their parents, we continue recursively backwards in time by treating the chosen mothers and fathers in the current stage as the offspring for the next stage. Using this sequential sampling scheme, we can compute *Pr*(*H|θ*), where *θ* = (*α, β, N*).

Furthermore, we can relate the mating parameters *α, β*, and *N* to the effective population size, *N*_*e*_, using the formula derived in (Gasbarra *et al.* 2005).

For the hyperprior, *P*(*θ*), we assume a uniform distribution for each of the parameters in *θ*. For instance, we assume *α* ~ *U*(*α*_*min*_, *α*_*max*_) for some fixed *α*_*min*_ and *α*_*max*_. We treat *β* and *N* in a similar way.

Finally, we combine the composite likelihood, prior, and hyperprior to approximate the joint posterior distribution of *H* and *θ* with

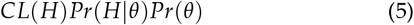

#### Markov Chain Monte Carlo

To explore the vast parameter space in a computationally feasible way, we use Markov-Chain Monte Carlo (MCMC) to sample from the posterior distribution of *H* and *θ*, approximated by Equation 5.

We represent the pedigree for a sample of individuals as a directed graph, where a node corresponds to an individual with a particular sex (i.e. male or female) and an edge represents a parent-offspring relationship. Individual *i* in the graph is not necessarily represented in the sample; but if it is sampled, the node is associated with a genotype vector *X*_*i*_. A more detailed description of the graph representation of pedigrees and the conditions for a valid pedigree is provided in File S1.

The MCMC explores pedigrees and mating parameters simultaneously. To explore the pedigree space, we make local modifications to the edges and the nodes in the graph using 10 reversible updates. The 10 updates can broadly be categorized into two groups. The first category of updates involves inserting or deleting edges to join or split pedigrees. The second category is modifying the pairwise relationship between two randomly chosen individuals, such as changing half-siblings to full-cousins, and vice versa. To explore the mating parameters, we use three different updates–one for each mating parameter–where we propose a new state by sampling the new parameter value from a normal distribution centered at the current value. A more detailed treatment of the updates is given in File S1.

Here, we outline the MCMC algorithm. Let *Q* = (*H, θ*) denote the set of parameters we want to estimate (i.e. pedigree and mating parameters).

1. Initialize pedigree *H* to be the one in which every individual is unrelated to each other. Initialize *α* by sampling from *U*(*α*_*min*_, *α*_*max*_), for some fixed *α*_*min*_ and *α*_*max*_. Initialize *β* and *N* in a similar way. Compute and store Equation 5 for the current configuration.
2. Choose one of the 10 updates at random and generate a new configuration.
3. If the new configuration is invalid, reject and go back to step 1. If it is valid, accept the new configuration with probability

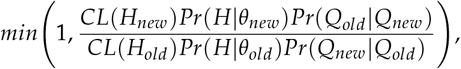
4. Repeat steps 1-3 *T* times.

The total number of samples, *T*, was chosen to achieve a balance between convergence of the Markov chain and computational time. Since we only want to keep samples after the Markov chain has converged to the stationary distribution, we discarded the first *B* samples as burn-in. To check for convergence, we ran multiple independent MCMC chains and checked that all chains fluctuated in a similar, stable range of log likelihood values. We note that this is only a proxy for checking convergence and there are other, albeit more involved, methods, such as checking the potential scale reduction factor for some specified quantity (Gelman *et al.* 1992). Furthermore, we keep only every *t*th sample to avoid storing correlated samples.

For both simulated and empirical datasets, which will be described next, we ran the MCMC for *T* = 6 × 10^6^ iterations with a burn-in period of *B* = 4 × 10^6^ iterations. The hyperprior for the mating parameters was set as follows: *α* ~ *U*(.1, 100), *β* ~ *U*(1 × 10^−5^, .1), and *N* ~ *U*(5, 5000). We also thinned the MCMC samples by keeping only every 50th sample.

### Simulated Data

We tested the performance of our method on simulated data. We simulated pedigrees up to two generations back in time using the mating model described in Prior with *α* = 15, *β* = 15, and *N* = 1000, which translates to *N*_*e*_ = 650 using the formula given in (Gasbarra *et al.* 2005).

We then simulated 10,000 independent single nucleotide polymorphic sites (SNPs) for each of the *N* founders in the pedigree, where the population allele frequency for each marker was sampled from the site frequency spectrum under neutrality. We assumed that the markers were spread evenly among 20 independent chromosomes of length 100Mb, and assumed sequencing error rate of .01. To test the effect of marker type on our parameter inference, we also simulated 20 microsatellites with 10 alleles of equal frequency per marker. Furthermore, we assumed that each marker was on an independent chromosome, with sequencing error rate of .01 and allele dropout rate of .05.

We then simulated the genotypes for the children in the pedigree by recombining parental haplotypes at rate 1.3e-8 per base pair per generation. We generated 50 independent datasets for both SNP and microsatellite simulations. For convenience, we refer to the simulations with SNPs as Simulation A and those with microsatellites as Simulation B in later sections.

### Empirical Data

We applied our method to reconstruct the previously unreported pedigree of house sparrows collected from an archipelago off the Helgeland coast of northern Norway (Lundregan *et al.* 2018). The individuals were genotyped using a custom Affymetrix 200K SNP array, with markers distributed across 29 of the chromosomes in the genome. Also provided were the location and year in which each each individual was collected.

We used individuals from a single island (island 27) to avoid any potential substructure in the sample. Furthermore, we restricted our analysis to the individuals born in 2009 to ensure that all samples belonged in a single generation. We pruned the markers for linkage disequilibrium (LD) using PLINK (Chang *et al.* 2015) at *r*^2^ = .05 to get a set of independent or loosely linked markers. The filtering steps resulted in 79 individuals and 4519 SNPs.

### Evaluation of Method

We compared the performance of our method to that of COLONY (Jones and Wang 2010), one of the most widely used pedigree reconstruction methods. We chose COLONY for several reasons. First, it supports full likelihood computation, which provides a gold standard to which we can compare our composite likelihood method. Second, it supports both SNPs and microsatellites data, allowing us to compare the performance of different marker types. Third, COLONY can estimate the short-term *N*_*e*_ based on the estimated frequency of siblings in the sample, which was shown to be more accurate than other methods of estimating *N*_*e*_ (Wang 2009).

Because the sample size in our simulations was much smaller than the population size, many pedigrees for the sample had similar likelihoods, making it difficult for both our method and COLONY to find the correct pedigree in its entirety. So we used pairwise prediction accuracy as a proxy for the accuracy of pedigree inference. In our method, we assigned pairwise relationship *R* to pair (*i, j*) if it had the highest posterior probability among all competing relationships. We approximated the posterior probability of *R* by counting the proportion of times pair (*i, j*) had relationship *R* in the MCMC samples. Similarly, we assigned relationship *R* to pair (*i, j*) in COLONY if it had the highest probability among all candidate relationships. Because the number of possible pedigrees is large, COLONY archives only the top *w* pedigrees with highest likelihoods. Suppose *S* is the set of indices for the pedigrees where (*i, j*) has relationship *R*. Then the probability of *R* is estimated by

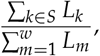

where *L*_*m*_ is the likelihood of the *m*th pedigree.

Furthermore, since COLONY restricts its inference to pedigrees going back only one generation back in time (i.e. siblings), we also limited our inference to the same scope when comparing the performance of our method to COLONY. The parameters used to run COLONY are detailed in File S2.

### Data Availability

Our software for pedigree inference is available for download at https://github.com/amyko/mcmcPed. Simulated data are available upon request. All supplemental files are available at FigShare.

## Results

### Simulated Datasets

To illustrate some of the issues involved in estimating multi-generation pedigrees, we first turn our attention to an example from Simulation A. Figure 1 shows the two most likely local pedigrees involving three sampled individuals (shaded) and their estimated posterior probabilities. In the first pedigree, individual 3 forms a full first cousin relationship with the other two individuals (1 and 2), as opposed to a half first cousin relationship as in the second pedigree. Here, the true pedigree is shown by the first pedigree (Figure 1A), which had the highest posterior probability.

**Figure 1.**
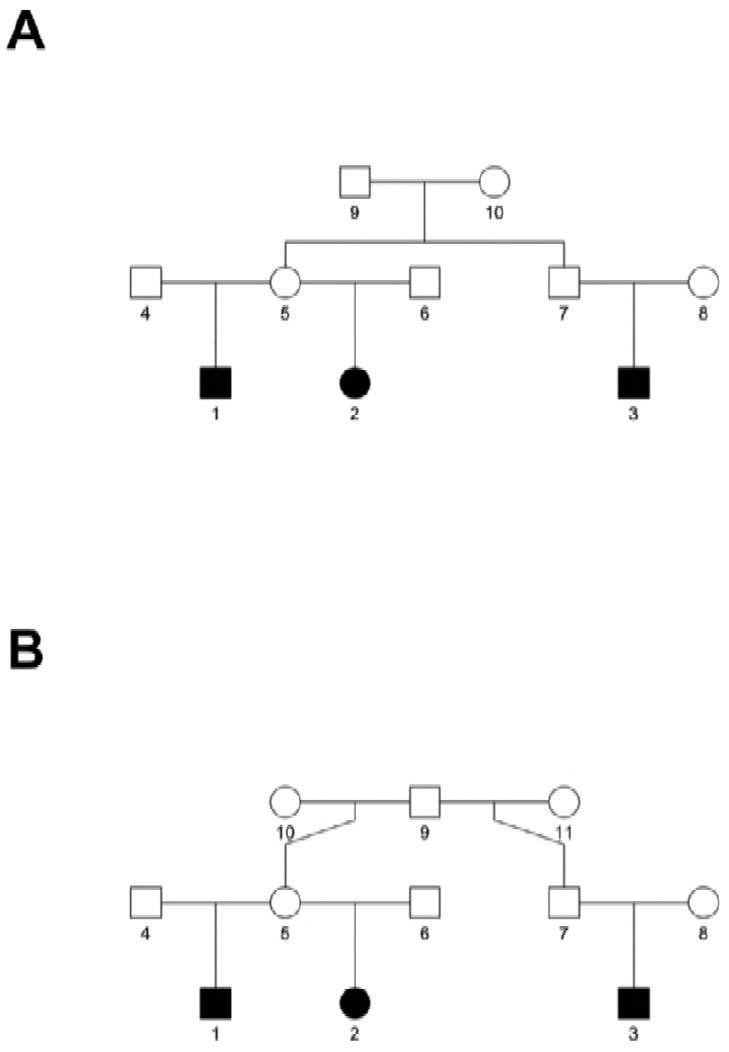
An example output pedigrees for three sampled individuals (shaded) from a dataset in Simulation A. Sex of the un-sampled individuals (unshaded) are unknown but are drawn in for illustration only. (A) Pedigree with the highest estimated posterior probability (*p* = .55). (B) Pedigree with the second highest estimated posterior probability (*p* = .45). The true pedigree is shown in panel A.

The uncertainty in the pedigree estimation, shown by the similar posterior probabilities of the two pedigrees (.55 and .45), was consistent with the fact that the pairwise likelihood values were similar under different relationships. More specifically, individuals 1 and 3 had a higher likelihood of being full cousins than half cousins by about one log likelihood unit. On the other hand, individuals 2 and 3 had a higher likelihood of being half cousins than full cousins by roughly the same amount. Based on pairwise likelihoods alone, individuals 1 and 3 would be classified as half cousins, and individuals 2 and 3 as full cousin. Piecing together such pairwise assignments, however, would not produce a valid pedigree. Such uncertainties in cousin inference were not uncommon: about 20 percent of true cousin pairs in Simulation A had nonzero posterior probabilities for both full and half cousins.

Table 1a shows the pairwise prediction accuracy of MCMC for the 50 independent datasets in Simulation A, where the pairwise likelihoods were computed using the method by (Albrechtsen *et al.* 2009). Full siblings, half siblings, and half cousins were classified correctly in almost all instances, whereas about seven percent of full cousin pairs were classified as half cousins. The rate of false detection of relatives was very low at about .01 percent, where the unrelated pairs were estimated as half cousins.

**Table 1.**
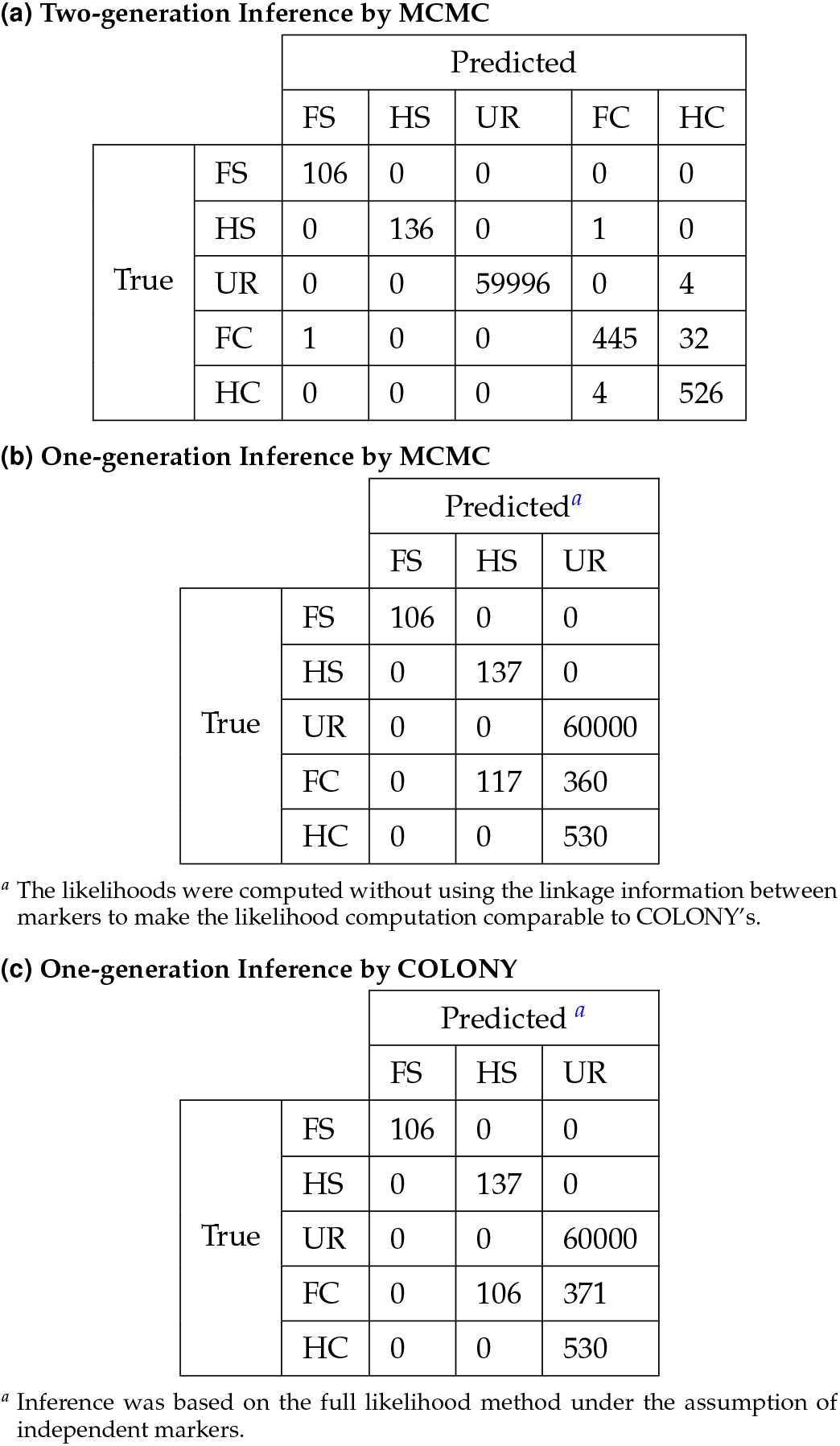
Pairwise Prediction Accuracy for Simulation A (SNPs)

**(a) Two-generation Inference by MCMC**
|  |  | Predicted |  |  |  |  |
| --- | --- | --- | --- | --- | --- | --- |
|  |  | FS | HS | UR | FC | HC |
| True | FS | 106 | 0 | 0 | 0 | 0 |
|  | HS | 0 | 136 | 0 | 1 | 0 |
|  | UR | 0 | 0 | 59996 | 0 | 4 |
|  | FC | 1 | 0 | 0 | 445 | 32 |
|  | HC | 0 | 0 | 0 | 4 | 526 |

**(b) One-generation Inference by MCMC**
|  |  | Predicted<sup>a</sup> |  |  |
| --- | --- | --- | --- | --- |
|  |  | FS | HS | UR |
| True | FS | 106 | 0 | 0 |
|  | HS | 0 | 137 | 0 |
|  | UR | 0 | 0 | 60000 |
|  | FC | 0 | 117 | 360 |
|  | HC | 0 | 0 | 530 |

<sup>a</sup> The likelihoods were computed without using the linkage information between markers to make the likelihood computation comparable to COLONY's.

**Table 1** Pairwise Prediction Accuracy for Simulation A (SNPs)
|  |  | Predicted<sup>a</sup> |  |  |
| --- | --- | --- | --- | --- |
|  |  | FS | HS | UR |
| True | FS | 106 | 0 | 0 |
|  | HS | 0 | 137 | 0 |
|  | UR | 0 | 0 | 60000 |
|  | FC | 0 | 106 | 371 |
|  | HC | 0 | 0 | 530 |

<sup>a</sup> Inference was based on the full likelihood method under the assumption of independent markers.

Figure 2A shows the posterior distribution of *N*_*e*_ estimated from the MCMC samples aggregated over the 50 datasets in Simulation A. The mode of the posterior distribution was close to the true value, indicated by the red vertical line. Similarly, Figure 2B shows that the distribution of maximum a posteriori (MAP) *N*_*e*_ for the 50 datasets was concentrated around the true value. The three mating parameters that make up the components terms of *N*_*e*_ (i.e. *α, β*, and *N*) showed high correlations among them. Figure S1 shows that high values of *N* tended to co-occur with low values of *α* for this simulation, which suggests that these parameters should not be estimated independently of each other and that marginal point estimates of any of these parameters are likely to be misleading.

**Figure 2.**
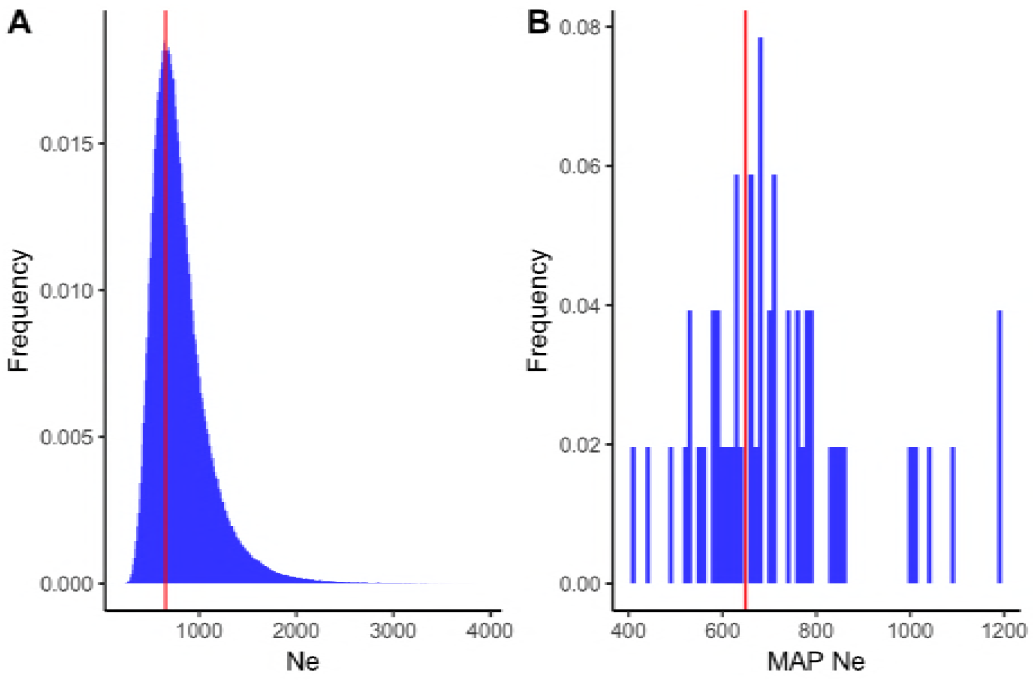
(A) Estimated posterior distribution of *Ne* from MCMC samples aggregated over 50 datasets in Simulation A. (B) Distribution of maximum a posterior (MAP) *N*_*e*_ for the 50 datasets in Simulation A. The red vertical line in each panel corresponds to the true value of the parameter.

Tables 1b and 1c compare the performance of our method with that of COLONY. Since COLONY estimates up to sibling relationships only, we also restricted the inference of our method to the same scope. Furthermore, we computed the likelihoods using the method discussed in (Weir *et al.* 2006) which assumes unlinked markers, an assumption that COLONY makes in its likelihood computation. Here, both our method and COLONY classified full siblings, half siblings, and unrelated pairs without error. Both methods also estimated all half cousin pairs to be unrelated. Furthermore, similar proportions of full cousin pairs were misclassified as half siblings by both methods: 22 percent by COLONY and 24 percent by our method. As shown in Figure 3, *N*_*e*_ was underestimated by both methods, which is consistent with the higher proportion of half siblings in the estimated pedigrees, caused by the misclassification of some full cousin pairs as half siblings.

**Figure 3.**
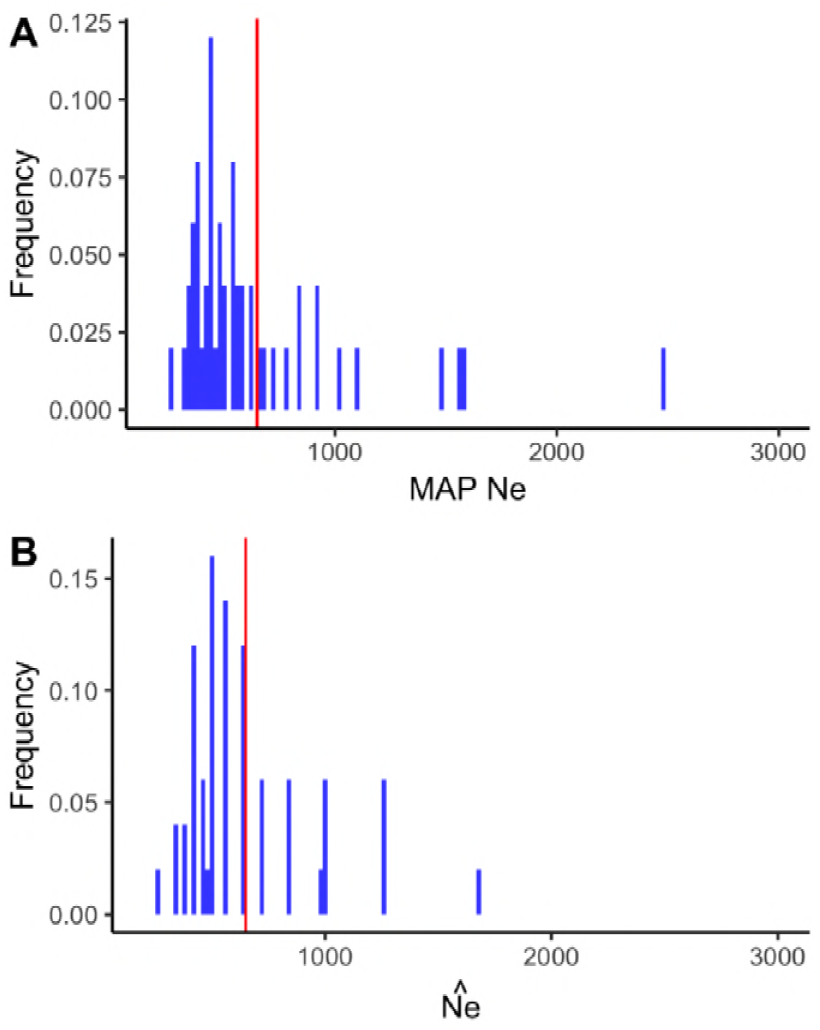
(A) Distribution of MAP *N*_*e*_ by MCMC, where the pedigree inference was restricted to one generation and the likelihood computation assumed independent markers. (B) Distribution of *N*_*e*_ estimates by COLONY based on full likelihood computation with independent markers and nonrandom mating.

Table 2a shows the pairwise prediction accuracy of MCMC for Simulation B (i.e. microsatellites), where the likelihoods were computed using the method by (Wang 2004). The accuracy rates were significantly lower than those in Simulation A (i.e. 10,000 SNPs). About 77 percent of full siblings and 27 percent of half siblings were classified correctly, and virtually all cousin pairs were estimated to be unrelated. This is likely due to the prior, which puts higher probabilities on sparsely connected pedigrees, overwhelming the likelihoods that do not show strong evidence for individuals being related. The distribution of MAP *N*_*e*_ also had a much higher variance compared to that of Simulation A (Figure 4A).

**Table 2.**
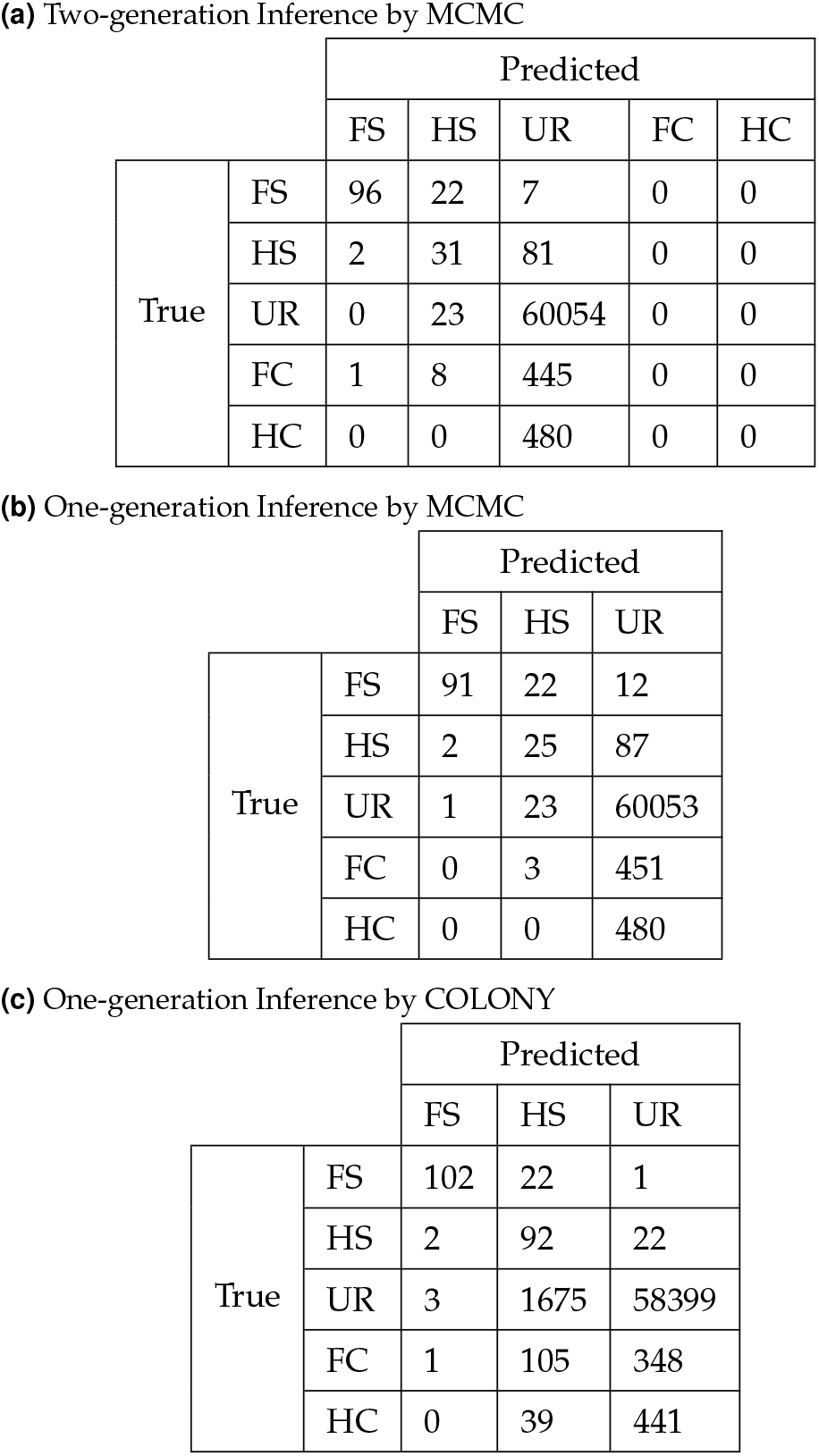
Pairwise Prediction Accuracy for Simulation B (Microsatellites)

**(a) Two-generation Inference by MCMC**
|  |  | Predicted |  |  |  |  |
| --- | --- | --- | --- | --- | --- | --- |
|  |  | FS | HS | UR | FC | HC |
| True | FS | 96 | 22 | 7 | 0 | 0 |
|  | HS | 2 | 31 | 81 | 0 | 0 |
|  | UR | 0 | 23 | 60054 | 0 | 0 |
|  | FC | 1 | 8 | 445 | 0 | 0 |
|  | HC | 0 | 0 | 480 | 0 | 0 |

**(b) One-generation Inference by MCMC**
|  |  | Predicted |  |  |
| --- | --- | --- | --- | --- |
|  |  | FS | HS | UR |
| True | FS | 91 | 22 | 12 |
|  | HS | 2 | 25 | 87 |
|  | UR | 1 | 23 | 60053 |
|  | FC | 0 | 3 | 451 |
|  | HC | 0 | 0 | 480 |

|  |  | Predicted |  |  |
| --- | --- | --- | --- | --- |
|  |  | FS | HS | UR |
| True | FS | 102 | 22 | 1 |
|  | HS | 2 | 92 | 22 |
|  | UR | 3 | 1675 | 58399 |
|  | FC | 1 | 105 | 348 |
|  | HC | 0 | 39 | 441 |

**Figure 4.**
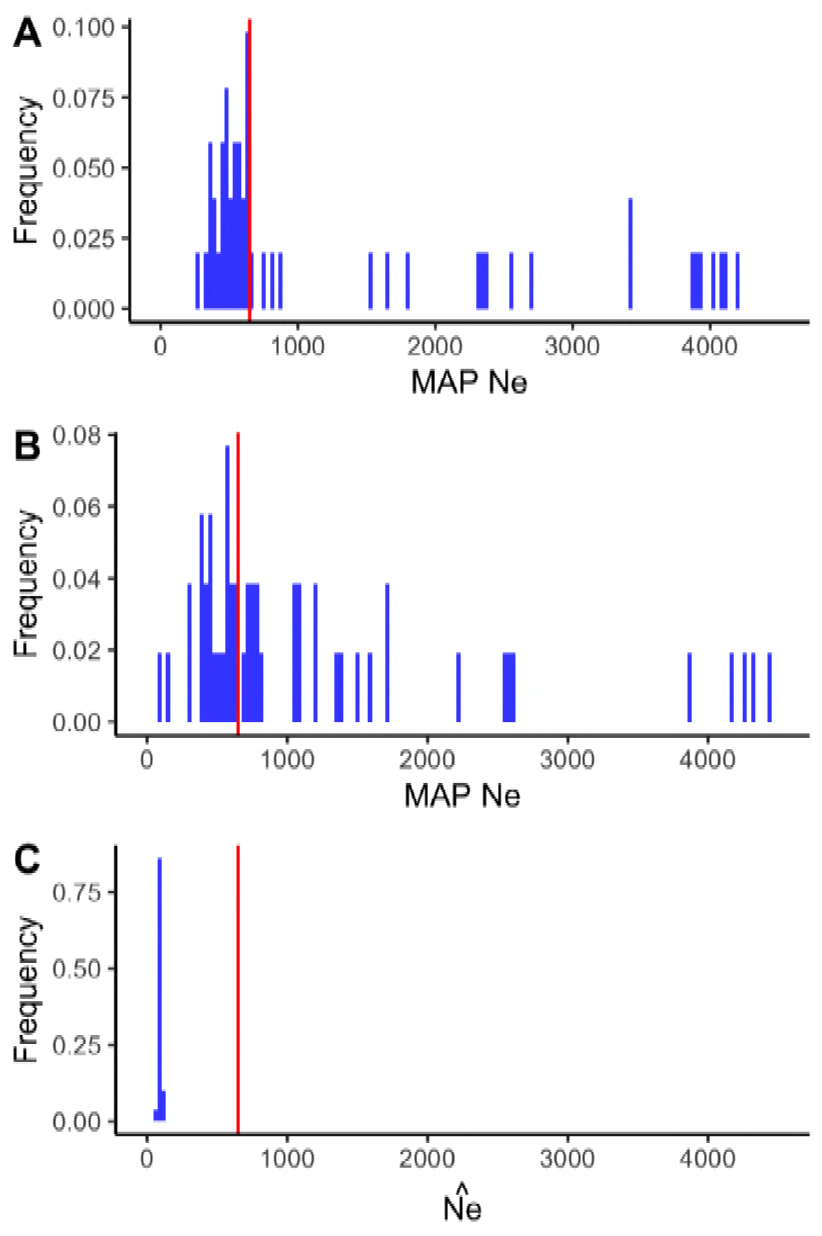
Distribution of the *N*_*e*_ estimates in Simulation B (i.e. microsatellites). (A) Distribution of MAP *N*_*e*_ estimated from MCMC samples under two-generation inference. (B) Distribution of MAP *N*_*e*_ estimated from MCMC samples under one-generation inference. (C) Distribution of *N*_*e*_ estimate by COLONY under nonrandom mating.

Tables 2b and 2c compare the performance of our method with that of COLONY for Simulation B. Again, we restricted the inference by our method to sibships to make a fair comparison with COLONY. Here, COLONY performed better than our method in correctly inferring full siblings and half siblings, but it also had a much higher false positive rate of 2.8 percent compared to .04 percent in our method. In fact, about 87 percent of the pairs estimated as half siblings by COLONY were actually unrelated. We note, however, that this problem may be addressed by adding an appropriate prior that is more conservative in half sibling assignments. Furthermore, due to the large number of unrelated pairs and cousins that were misclassified as half siblings, *N*_*e*_ was significantly underestimated by COLONY (Figure 4C).

For all the experiments, we checked the convergence of MCMC by studying the likelihood trace of multiple independent chains. As an illustration, we show an example of the log likelihood trace for the last one million iterations for a single experiment in Simulation A (Figure S2).

The running time for our method depends on many factors, such as the sample size, the underlying pedigree structure, and the maximum number of generation allowed in the pedigree inference. As an example, an MCMC run with 6 million iterations for a two-generation pedigree inference took about 36 seconds on a laptop with 2.3 GHz Intel Core i5 processor for a single dataset in Simulation A, excluding the pre-computation time for calculating the likelihoods.

### Effect of Presence of Relatives Beyond First Cousins

For real datasets, it is often unreasonable to assume that the sample does not contain relatives more distant than first cousins. Here we show the effect of having second cousins in the sample on the inference of pedigrees and *N*_*e*_. Table 3 shows the prediction accuracy for a simulation scenario where second cousins were present in the sample. The simulation parameters were identical to those of Simulation A, except for the number of generations under which the pedigrees were simulated. Instead of going back up to two generations back in time as in Simulation A–which generated relatives up to first cousins–here we simulated pedigrees up to three generations back in time, which generated second cousins as well.

**Table 3.** Pairwise Prediction Accuracy for Datasets Containing Second Cousins (Inference by MCMC)

|  |  | Predicted |  |  |  |  |
| --- | --- | --- | --- | --- | --- | --- |
|  |  | FS | HS | UR | FC | HC |
| True | FS | 118 | 1 | 0 | 0 | 0 |
|  | HS | 0 | 108 | 2 | 0 | 1 |
|  | UR | 0 | 0 | 56189 | 0 | 3 |
|  | FC | 5 | 5 | 0 | 386 | 95 |
|  | HC | 0 | 0 | 9 | 4 | 499 |
|  | 2FC | 0 | 0 | 523 | 2 | 1388 |
|  | 2HC | 0 | 0 | 1482 | 0 | 430 |

As we can see in Table 3, the accuracy rates were similar to those of Simulation A for relationships up to first cousins. However, about 73 percent of full second cousins (2FC) were classified as half first cousins (HC), the most distant relationship type our method is designed to estimate. Similarly, about 22 percent of half second cousins (2HC) were classified as HC. As expected, *N*_*e*_ was biased downward due to the high frequency of HC in the estimated pedigrees, caused by the misclassification of second cousins as HC (Figure 5).

**Figure 5.**
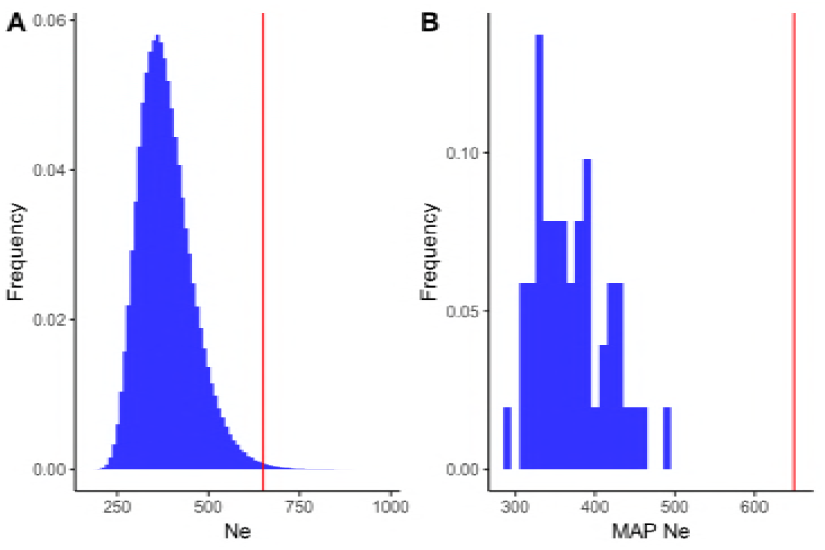
(A) Estimated posterior distribution of *Ne* from MCMC samples aggregated over 50 datasets, where the data contained second cousins. (B) Distribution of MAP *N*_*e*_ for the 50 datasets.

To correct the downward bias in *N*_*e*_ estimation, we took advantage of the fact that our method can still infer siblings with high accuracy (Table 3). More specifically, we simulated pedigrees under various *N*_*e*_ to find a value that generated a number of siblings close to the one estimated by our method. Let *S*_*IBD*_ = *N*_*FS*_ + .5*N*_*HS*_ be the summary statistic that measures the level of identical-by-descent (IBD) contributed by siblings in the sample, where *N*_*FS*_ and *N*_*HS*_ are the number of full siblings and half siblings, respectively; and denote *Ŝ*_*IBD*_ to be the statistic obtained from the MCMC inference on the sample. Let *α*_*MAP*_ and *β*_*MAP*_ be the MAP estimates of *α* and *β*, respectively, computed using the marginal posterior distributions obtained from the MCMC samples. We then simulated pedigrees going back up to one generation in time under *α*_*MAP*_, *β*_*MAP*_, and various values of *N*–which translates to different values of *N*_*e*_–and computed *S*_*IBD*_ from the simulated pedigrees. We then chose the value of *N*_*e*_ that produced *S*_*IBD*_ that most closely matched *Ŝ*_*IBD*_.

Figure 6 shows the distribution of the *N*_*e*_ estimates after correcting for bias as described above. Although the standard error was higher than that of uncorrected estimates, the median of the distribution (657) was much closer to the true value (650) than before.

**Figure 6.**
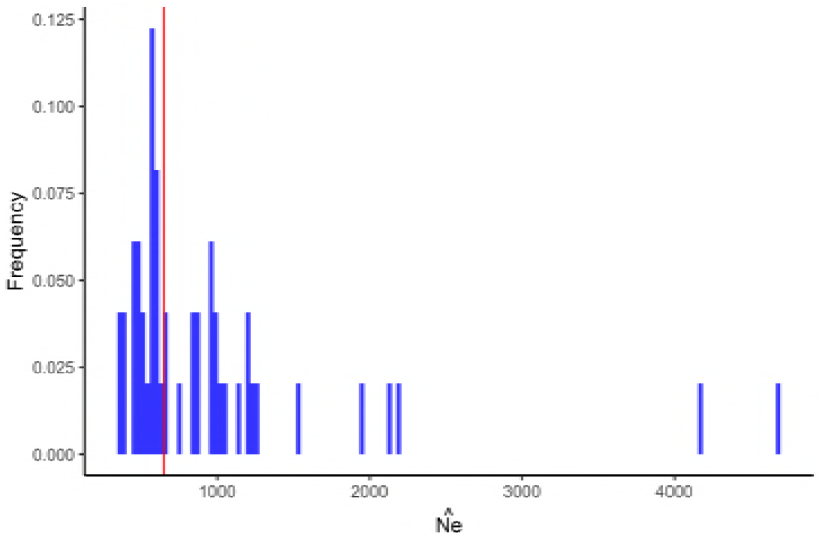
Distribution of the *N*_*e*_ estimates for the 50 datasets after bias correction. The red vertical line indicates the true value of *N*_*e*_.

### Sparrow Dataset

We analyzed a subset of the house sparrow dataset sequenced by Lundregan *et al.* (2018). After the filtering steps described in Empirical Data, the sample consisted of 75 individuals and 4,519 SNPs distributed across 29 autosomes. Here we show an example of the inferred pedigrees by our method and compare them to those estimated by COLONY.

Figure 7 shows the likely local pedigrees involving five individuals (shaded) in the sparrow dataset. The estimated posterior probabilities of the pedigrees shown in panel A and B were .77 and .23, respectively. The difference between the two pedigrees was the pairwise relationship between individuals 1339 and 1450, which was estimated to be full cousins in panel A and half cousins in panel B. Figure 7C shows the pedigree with the highest likelihood estimated by COLONY. This pedigree had posterior probability of zero in our method. We see that the half sibling relationship between individuals 1390 and 1450 were recovered by COLONY but all cousin relationships that our method detected were estimated to be unrelated. Based on the simulation studies in Simulated Datasets, however, we expect the full first cousin relationships inferred by our method to be either true first cousins or, with considerably smaller probability, more distant relatives (e.g. second cousins).

**Figure 7.**
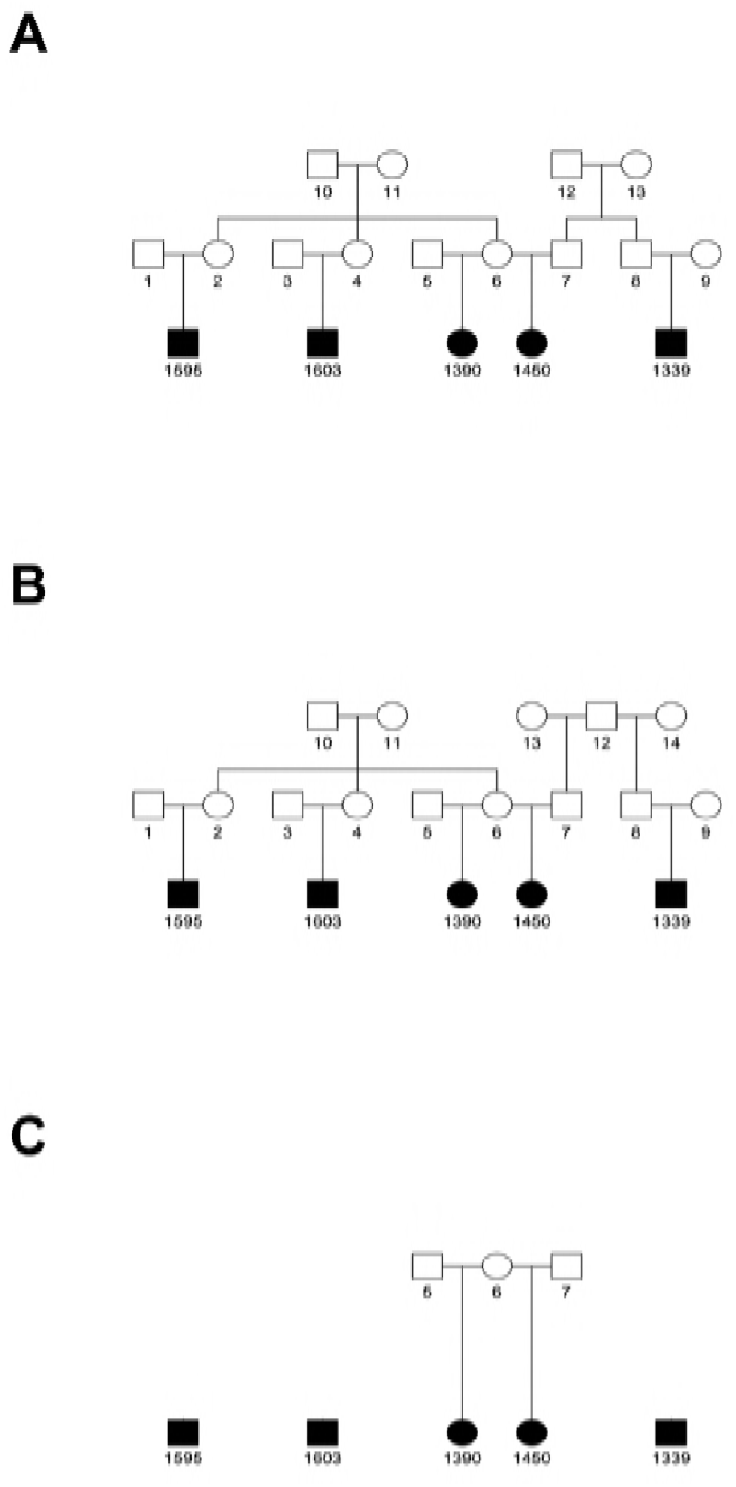
Estimated pedigrees of five sampled individuals in the sparrow dataset. (A) Pedigree with estimated posterior probability of .77. (B) Pedigree with estimated posterior probability of .23. (C) Most likely pedigree estimated by COLONY, but whose posterior probability was zero in our method.

Table 4 compares the pairwise relationship classifications between our method and COLONY. Pairs that were classified as full siblings, half siblings, or unrelated by our method largely agreed with the classifications by COLONY. On the other hand, about 29 percent of pairs that were estimated to be full cousins by our method were estimated to be half siblings by COLONY, which is consistent with what was observed in the simulation studies in Simulated Datasets. Furthermore, most of the relationships that were inferred as half cousins by our method were classified as unrelated by COLONY.

**Table 4.**
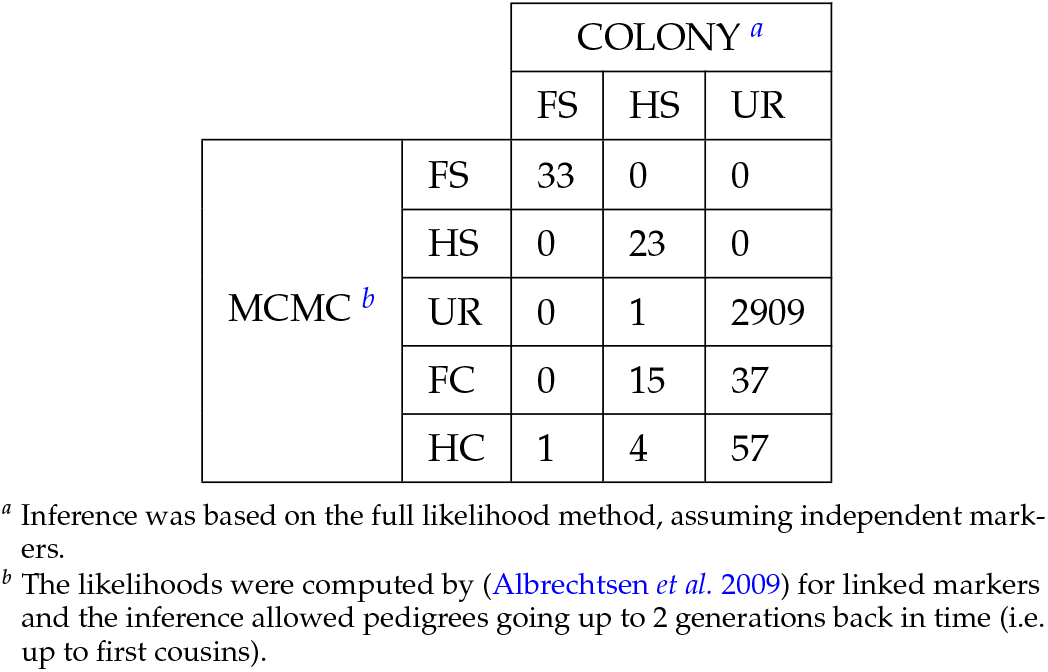
Comparison of Pairwise Relationship Classification by MCMC and COLONY.

|  |  | COLONY <sup>a</sup> |  |  |
| --- | --- | --- | --- | --- |
|  |  | FS | HS | UR |
| MCMC <sup>b</sup> | FS | 33 | 0 | 0 |
|  | HS | 0 | 23 | 0 |
|  | UR | 0 | 1 | 2909 |
|  | FC | 0 | 15 | 37 |
|  | HC | 1 | 4 | 57 |

<sup>a</sup> Inference was based on the full likelihood method, assuming independent markers.
<sup>b</sup> The likelihoods were computed by (Albrechtsen et al. 2009) for linked markers and the inference allowed pedigrees going up to 2 generations back in time (i.e. up to first cousins).

## Discussion

We have shown that, given enough marker information, our method is able to jointly estimate *N*_*e*_ and relationships up to first cousins accurately and efficiently. Unlike existing pedigree inference methods, our method not only allows estimation of pedigrees and *N*_*e*_, but also provides an uncertainty measure on the estimates via posterior probabilities. Furthermore, our method provides a framework for incorporating different types of population models in the prior for the pedigrees, which can potentially allow us to estimate other population parameters, such as migration rates between subpopulations.

Our method also improves upon one of the most widely used pedigree reconstruction programs, COLONY, by estimating relationships beyond sibships. This not only expands the types of pedigrees we can infer but also increases the accuracy of sibship inference. In particular, first cousins were often misclassified as half siblings if the estimation method did not allow inference of cousins. For example, about 44 percent of half siblings estimated by COLONY using 10,000 SNPs were actually first cousins (Table 1c). Furthermore, we showed that *N*_*e*_ can be underestimated if the sample contains cousins but the pedigree inference is restricted to sibships only (Figure 3). By explicitly including first cousins in the inference, our method was able to infer half siblings with higher precision (Table 1a), as well as estimate *N*_*e*_ more accurately (Figure 2). However, we note that the problem persists when the sample contains relatives more distant than first cousins. When datasets contained second cousins, for example, they were often estimated as half first cousins–the most distant relationship our method is designed to estimate–and consequently caused a downward bias in *N*_*e*_ estimates. Therefore, we must use caution in interpreting inferred half cousins, as the true relationship could be more distant, and use the simulation method discussed in Effect of Presence of Relatives Beyond First Cousins to correct for potential bias in *N*_*e*_ estimates.

We note that the performance of our method relies heavily on the accuracy of pairwise likelihoods. The accuracy of pairwise likelihoods depends on many factors, such as marker density, level of linkage disequilibrium, sequencing error rates, and population allele frequency estimates. Ignoring the linkage between markers, in particular, significantly decreased the power to detect first cousins (File S3). Due to linkage, close relatives such as first cousins are expected to share, with high probability, long IBD segments that are on the order of megabases in length, although the probability of IBD per marker is relatively low (Chapman and Thompson 2003). The presence of such long IBD segments should make detecting relatives quite easy even though identifying the exact relationship can be more difficult. Treating the markers as independent, however, does not take advantage of the presence of long IBD segments and thus decreases our ability to detect relatives (Tables 1b, 1c). Therefore, likelihood computation methods, such as (Albrechtsen *et al.* 2009), that take into account the linkage information between markers should be used instead for detecting relatives, and naturally, for pedigree inference as well.

Marker type and density also have a significant impact on the quality of pairwise likelihoods. We have seen that using 20 highly informative microsatellites performed worse than using 10,000 SNPs. The accuracy rates of COLONY (Table 2c) suggest that the use of microsatellites to estimate sibships might be misguided in practice since first cousins can often be misclassified as half siblings in methods that do not explicitly model first cousins. Furthermore, microsatellites may not provide enough information to easily distinguish between full and half siblings (Table 2). Also, 20 microsatellites with 10 alleles of equal frequency in our simulations is more generous than what is available in many real datasets, and the performance on less informative datasets is likely to be worse than what was shown in this study. We note that finding the best ways to address the various challenges in pairwise likelihood computation is an active area of research and requires further investigation.

There are limitations to our method that require further work. Our method does not support pedigrees that contain cycles, except those caused by full sibling relationships. More specifically, we do not consider pedigrees that are inbred or have complex, cyclic relationships such as double first cousins. A simulation study by (Ko and Nielsen 2017) suggests that in the presence of inbred individuals, the method will tend to estimate individuals to be more genealogically closer than they actually are (e.g. first cousins estimated half siblings). Furthermore, our method assumes that all samples belong in a single generation, which may not typically be true for many real datasets. This may be addressed by adding updates in the MCMC that allow sampled individuals to move between generations. Furthermore, our method does not yet scale up to sample sizes typical of GWAS as the number of pairwise comparisons still increases rapidly with sample size. One possible approach to address this issue is partitioning the sample into smaller sets using methods such as (Manichaikul *et al.* 2010) and estimating the pedigrees for each smaller subset of individuals.

Overall, our method provides a way to jointly estimate pedigrees and *N*_*e*_, and measure the uncertainty of the estimates in a computationally efficient way. Importantly, our method also provides a basic framework for estimating demographic parameters of the current population from pedigrees–analogous to population genetic methods based on coalescent trees–thus opening up new possibilities for learning about the demographic history of the recent past.

